# FLASH or flare: variable intestinal toxicity results in a mouse model following proton pencil beam scanning irradiation on a clinical superconducting synchrocyclotron

**DOI:** 10.1101/2025.09.17.676536

**Authors:** Eva Bogaerts, Ellina Macaeva, Sima Qamhiyeh, Laurence Delombaerde, Brigitte Reniers, Marco Caprioli, Nicolas Gerard, Jarrick Nys, Valentin Hamaide, Alexis Warnier, Rudi Labarbe, Swati Girdhani, Richard Coos, Stéphane Lucas, Sofie Isebaert, Rüveyda Dok, Evelien Dierick, Karin Haustermans, Edmond Sterpin

## Abstract

**Background and aims:** Ultra-high dose rate (FLASH) irradiation is a promising technique to reduce radiation-induced normal tissue toxicities while preserving antitumor efficacy. We evaluated the feasibility and intestinal sparing potential of FLASH irradiation using a clinical synchrocyclotron-based proton therapy system generating a pulsed beam.

**Material and methods:** C57BL/6J mice received abdominal irradiation (2×2 cm) in transmission mode at FLASH (>60 Gy/s) or conventional (CONV, 0.5 Gy/s) dose rates using a 230 MeV superconducting synchrocyclotron proton pencil beam scanning (PBS) system. Two independent irradiation rounds were performed. Endpoints included 75-day survival, regenerating crypt counts, whole blood counts at day 4, and intestinal wall thickness, cyst-like structures, and cytokine levels at day 75.

**Results:** In the first irradiation round, survival after 14.5 Gy FLASH was markedly improved (5/8 survivors) compared to CONV (0/8), whereas in the second round, survival rates were identical (2/7 per group). Overall, pooled data indicated improved survival with 14.5 Gy FLASH. The LD50 was 13.74 Gy in CONV and 14.48 Gy in FLASH mode, corresponding to a FLASH modifying factor of 0.95. FLASH at 14.5 Gy increased regenerating crypt numbers compared to CONV, but only in the first round, supporting survival outcomes. No significant differences were observed in whole blood counts, cytokine profiles, or long-term intestinal structural changes between groups.

**Conclusion:** FLASH proton therapy delivered with a clinical synchrocyclotron PBS system can reduce short-term gastrointestinal toxicity in mice. However, inconsistent results across irradiation rounds highlight limitations of this model for reliable FLASH studies.

## Introduction

FLASH radiotherapy (RT), an innovative technique delivering radiation at ultra-high dose rates (UHDR; >40 Gy/s), has re-emerged in the RT field. First demonstrated in 2014, FLASH RT reduced lung toxicity compared to conventional (CONV) irradiation while maintaining antitumor efficacy [1] —a differential effect now known as the FLASH effect.

Several research groups have already been able to observe the FLASH normal tissue sparing and/or iso-effective tumor eradication in various animal models with various radiation sources [2-8]. Early findings mainly relied on low-energy electrons with limited penetration, restricting clinical use. In contrast, protons offer deep-tumor treatment with superior dose distribution and sparing of organs at risk.

Clinically, the first proton FLASH trial (FAST-01, 2023) confirmed feasibility and safety in patients with extremity bone metastases [9]. A second proton FLASH clinical trial enrolling patients with painful thoracic bone metastases (FAST-02) is ongoing [10]. Despite the fast translation of this experimental treatment modality, solid evidence of the biological mechanisms behind the FLASH effect is lacking. Moreover, negative pre-clinical results have also been reported. Bell and colleagues found that proton FLASH (≥80 Gy/s) whole abdominal irradiation significantly increased mortality and induced similar changes in intestinal histology compared to CONV irradiation [11]. Additionally, a recent study involving proton FLASH (100 Gy/s) partial abdominal irradiation could not demonstrate any tissue-sparing effect [12].

Most *in vivo* proton FLASH studies used quasi-continuous cyclotron beams [13-20]. In this proof-of-concept preclinical mouse study, we assessed the technical feasibility and intestinal sparing potential of 230 MeV S2C2 clinical proton pencil beam scanning (PBS) FLASH irradiation.

## Materials and methods

### S2C2 beam time structure

The IBA S2C2 synchrocyclotron accelerates one pulse of protons every millisecond (Pulse Repetition Frequency, *PRF* = 1 kHz) [21]. As the pulse lasts for 7 μs (*t*_pulse_), there are 993 μs between subsequent pulses (Δ*t*_pulse_). The beam intensity is modulated only by means of charge per pulse (*Q*_pulse_) modification, which in turn is modulated through the average radiofrequency system amplitude during the whole acceleration cycle. Schematic representation of the S2C2 beam time structure is shown in Supplementary Figures 1 and 2.

### Treatment plan and beam delivery

Pencil Beam Scanning technique was used to deliver the shoot through beam. Two treatment plans were designed for CONV and UHDR irradiations respectively (Figures 1A and 1C). A custom-made brass aperture (20×20 mm) was used to constrain the lateral profile. The total number of MUs was determined by measurement of the central dose during the dosimetry campaign as described below in Dosimetry section. Further details can be found in Supplementary material.

**Figure 1.**
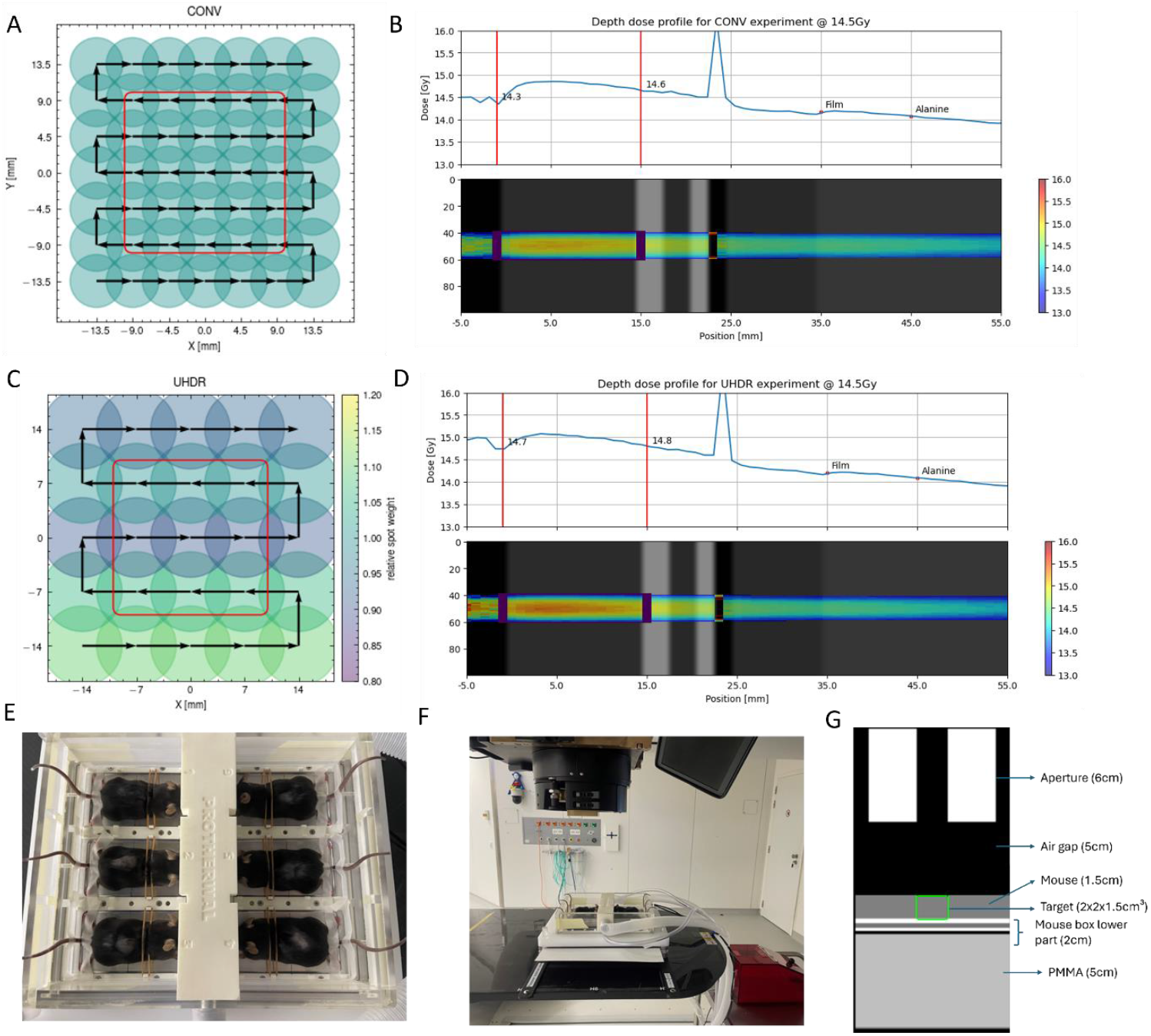
Treatment planning and irradiation setup. Single energy PBS pattern for CONV (A) and UHDR (C) irradiation. In color, the relative spot weight. Arrows indicate the scanning pattern. The aperture opening is projected in the isocenter plane (red square). The spot spacing equals approximately 1.3 times the average spot size (σ_X_ + σ_Y_)/2. Dose deposition across depth for CONV (B) and UHDR (D) experiments at 14.5Gy on the in-silico model. The depth-dose curve is averaged over 10 mm laterally. The two vertical red lines delimit the mouse (modeled as 1.5 cm of water). Downstream to the mouse are the IRRAMICE box, the gafchromic film and the alanine pellet. (E) Custom IRRAMICE anesthesia and positioning box, fitting six mice in individual compartments. (F) IRRAMICE box on top of the dosimetric set-up on the patient couch in the clinical treatment room. (G) Schematic representation of the irradiation setup used for *in silico* modelling. From upstream to downstream, we have: 6cm brass aperture, 5cm air gap, 1.5cm water (representing mouse), 2cm of mice box holder consisting of different materials (aluminium, polyurethane, air, high density polyethylene), >5cm PMMA.

### In silico

An *in silico* model of the entire experimental setup (custom-made brass aperture, mice, animal positioning box) (Figure 1G) was developed to study dose profile across depth inside the mice via Monte Carlo (MC) simulations. The *in silico* model was validated with the film (EBT-XD) and alanine pellet measurements. Further details can be found in Supplementary material.

### Dosimetry

The monitor unit-to-dose relation was established in a dosimetry session in which Gafchromic EBT-XD films (AshLand Inc.), and alanine pellets (Harwell Inc.) were placed between layers of PMMA and irradiated simultaneously. The central average dose (0.5×0.5 cm) per field was extracted from the EBT-XD films and a linear fit was performed to relate monitor unit to dose. EBT-XD films were calibrated prior using conventional proton beams with the two-page calibration protocol by Crijns et al. [22]. The alanine pellets were used for independent verification. The dose homogeneity throughout the treatment volume was assessed using the EBT-XD film.

### Animals

Eleven-week-old female C57BL/6J mice were acquired from Janvier Labs (France). Table 1 summarizes all experimental endpoints, doses and numbers of animals used. Doses ranging between 12.5 and 15.5 Gy in FLASH mode and between 13 and 16 Gy in CONV mode were used to calculate the LD50. For the rest of the endpoints, 13.5 and 14.5 Gy were used in both modes. Irradiations were evenly distributed across two weekends separated by several months, with CONV treatments conducted on Friday afternoons and FLASH treatments on Saturday afternoons. All experimental procedures were approved by the Ethical Committee for Animal Experimentation (ECD) of KU Leuven (P172/2021 and P133/2024) in compliance with national and European regulations. Further details can be found in Supplementary material.

**Table 1.**

| **Dose, Gy** | **Mode** | **Nr of mice included in each irradiation round (1 or 2) \*** | **Endpoints** |
| --- | --- | --- | --- |
| 12.5 | FLASH | 6 mice in round 2 | Survival |
| 13.5 | FLASH | 8 mice in round 1<br>7 mice in round 2 | Survival, cytokine analysis and histology at 75 days post-irradiation |
| 13.5 | CONV | 8 mice in round 1<br>7 mice in round 2 | Survival, cytokine analysis and histology at 75 days post-irradiation |
| 13.5 | FLASH | 4 mice in round 1<br>5 mice in round 2 | Crypt survival assay, whole blood counts and cytokine analysis at 4 days post-irradiation |
| 13.5 | CONV | 4 mice in round 1<br>5 mice in round 2 | Crypt survival assay, whole blood counts and cytokine analysis at 4 days post-irradiation |
| 14.5 | FLASH | 8 mice in round 1<br>7 mice in round 2 | Survival, cytokine analysis and histology at 75 days post-irradiation |
| 14.5 | CONV | 8 mice in round 1<br>7 mice in round 2 | Survival, cytokine analysis and histology at 75 days post-irradiation |
| 14.5 | FLASH | 4 mice in round 1<br>5 mice in round 2 | Crypt survival assay, whole blood counts and cytokine analysis at 4 days post-irradiation |
| 14.5 | CONV | 4 mice in round 1<br>5 mice in round 2 | Crypt survival assay, whole blood counts and cytokine analysis at 4 days post-irradiation |
| 15.5 | FLASH | 6 mice in round 2 | Survival |
| 14.0 | CONV | 6 mice in round 1 | Survival |
| 13.0 | CONV | 6 mice in round 1 | Survival |
| 16.0 | CONV | 6 mice in round 1 | Survival |

\*The number of samples shown in the graphs may vary due to the loss of animals from causes unrelated to the treatment and sample loss during processing.

### Animal positioning

For abdominal irradiation (2×2 cm collimated field), mice were placed in a custom IRRAMICE anesthesia and positioning box [23] fitting six mice in individual compartments (Figures 1E and1F). Isoflurane (2% in medical air) was used to anesthetize mice during the irradiation treatment. Further details can be found in Supplementary material.

### EdU staining

Mice were injected intraperitoneally with 200 µg of 5-ethynyl-2’-deoxyuridine (EdU; Invitrogen; #E10044) on day 4 post-irradiation and sacrificed 3 hours post-injection. Intestinal segments were collected by the Swiss-roll technique, and processed as described previously [24].

Seven micrometer thick sections (at least three nonadjacent sections/sample) were labeled using the Click-iT EdU Alexa Fluor 488 Imaging Kit (Invitrogen; #C10337) according to the manufacturer’s instructions. The number of regenerating crypts (≥5 EdU+ cells/crypt) was counted in the most severely damaged region. Further details can be found in Supplementary material.

### Histology

Surviving animals at 75 days post-irradiation were euthanized and intestinal segments were collected by the Swiss-roll technique as described above. Five micrometer thick sections were cut and stained with hematoxylin and eosin. The mean thickness of the intestinal wall and the average number of intra-mucosal cyst-like structures were quantified. Masson Trichrome kit (Biognost; #MST-K-500) was used to qualitatively check for collagen deposition. Further details can be found in Supplementary material.

### Complete blood count analysis

Whole blood from the posterior vena cava was collected into EDTA-coated tubes at four days post-irradiation. Complete blood count data were obtained by analyzing the blood using a scil Vet abc Plus (Scil Animal Care).

### Cytokine quantification

Blood was collected on day 4 and 75 post-irradiation from the posterior vena cava using a 26G needle and an EDTA-coated syringe. Plasma was obtained by centrifugation of whole blood for 10 min at 1,000 x g using a refrigerated centrifuge. Plasma samples were stored at -80°C. Cytokine concentrations were quantified using an MSD U-PLEX electro-chemiluminescent immunoassay (Meso Scale Diagnostics) following the manufacturer’s instructions. Biomarker assays used were custom 96-well 6-plex plates for simultaneous analysis of species-specific IFNγ, IL6, IL10, KC-GRO/CXCL-1, MCP-1 and TNFα (K15069M-1) and 96-well single-plex plates for the analysis of TGFβ1 (K152ADM-1).

### Statistical analysis

Statistical analyses were performed with GraphPad Prism 10.2.3. Relative body weights were compared with multiple t-tests, survival with the Log-rank (Mantel-Cox) test, and regenerating crypt numbers with t-tests. Intestinal wall thickness, cyst counts, and blood counts were analyzed with Kruskal-Wallis and Dunn’s post hoc tests. Cytokine levels were assessed with one-way ANOVA with Tukey’s post hoc test (day 4) or Kruskal-Wallis test with Dunn’s post hoc test (day 75). Survival versus dose was modeled by simple linear regression. p<0.05 was considered significant.

## Results

### In silico model for dose and dose rate calculation

A Monte Carlo computation was performed on the *in silico* model described in the Materials and Methods section from the nozzle exit up until the bottom of the IRRAMICE box.

The depth dose profile is depicted in Figures 1B and 1D for the experiments at CONV and UHDR with a target dose of 14.5 Gy. The dose deposition profile was relatively flat across the depth inside the mouse body. However, the dose decreased laterally from the center of the field to the aperture edges. This was confirmed on the EBT-XD film measurements. For the CONV irradiation, the mice received a median dose of 14.03 Gy, and for the UHDR irradiation, a median dose of 14.2 Gy on the 2×2×1.5cm^3^ target volume while the median doses are 14.80 Gy and 15.09 Gy on a central region of 1×1×1.5 cm^3^ for the CONV and UHDR respectively. The full dose-volume histograms (DVHs) for the CONV and UHDR irradiation are available in Supplementary Figure 3.

For 14.5 Gy CONV irradiation, median dose variations were <1% with alanine pellets and 3– 6% with films compared to simulations. For the same dose at UHDR, variations were <1% with alanine pellets and 1.5–6% with films (Supplementary Tables 1 and 2).

A dose delivery simulation was also performed with a maximum of 6.6 MUs per pulse. This led to the delivery of the UHDR field in 220 ms, which is the same as the delivery time during the first round of UHDR irradiation at 14.5 Gy. The resulting dose-rate volume histogram is available in Supplementary Figure 4. According to the dose rate definition used in the max-percentile dose rate at the 95th percentile [25], 95% of the target volume received a dose rate of at least 74 Gy/s. This is slightly higher than the average field dose rate computed as 14.5 Gy/220 ms=66 Gy/s.

### FLASH proton irradiation results in reduced mortality compared to CONV proton irradiation, however, different results are obtained across independent irradiation rounds

To evaluate whether FLASH proton irradiation reduces morbidity and mortality compared to CONV, mouse body weight and survival were monitored for 75 days (Figure 3). At 14.5 Gy, FLASH-treated mice lost significantly less weight on days 3–5 post-irradiation than CONV-treated mice, though no other significant differences in weight loss were observed (Figure 3A). Survival at 13.5 Gy was similar between groups (13/15 FLASH vs. 12/15 CONV), but at 14.5 Gy survival was higher with FLASH (7/15) than CONV (2/15), suggesting a dose-dependent effect (Figure 3B). No mice survived higher dose (15.5 Gy FLASH, 16 Gy CONV) irradiations. All the animals surviving beyond 9 days after irradiation survived until the end of the study. Importantly, outcomes at 14.5 Gy differed between rounds (Figures 2D and2E): in round 1, FLASH mice showed reduced weight loss and 5/8 survived long-term, while all CONV mice required euthanasia by day 8; in round 2, in contrast, survival was low in both groups (2/7). Simple linear regression was used to determine the dose that led to 50% lethality (LD50) based on the pooled data of two irradiation rounds to obtain enough data points for curve fitting (Figure 3C). The CONV LD50 was determined to be 13.74 Gy and the FLASH LD50 was 14.48 Gy, which corresponds to a FLASH modifying factor (FMF) of 0.95.

**Figure 2.**
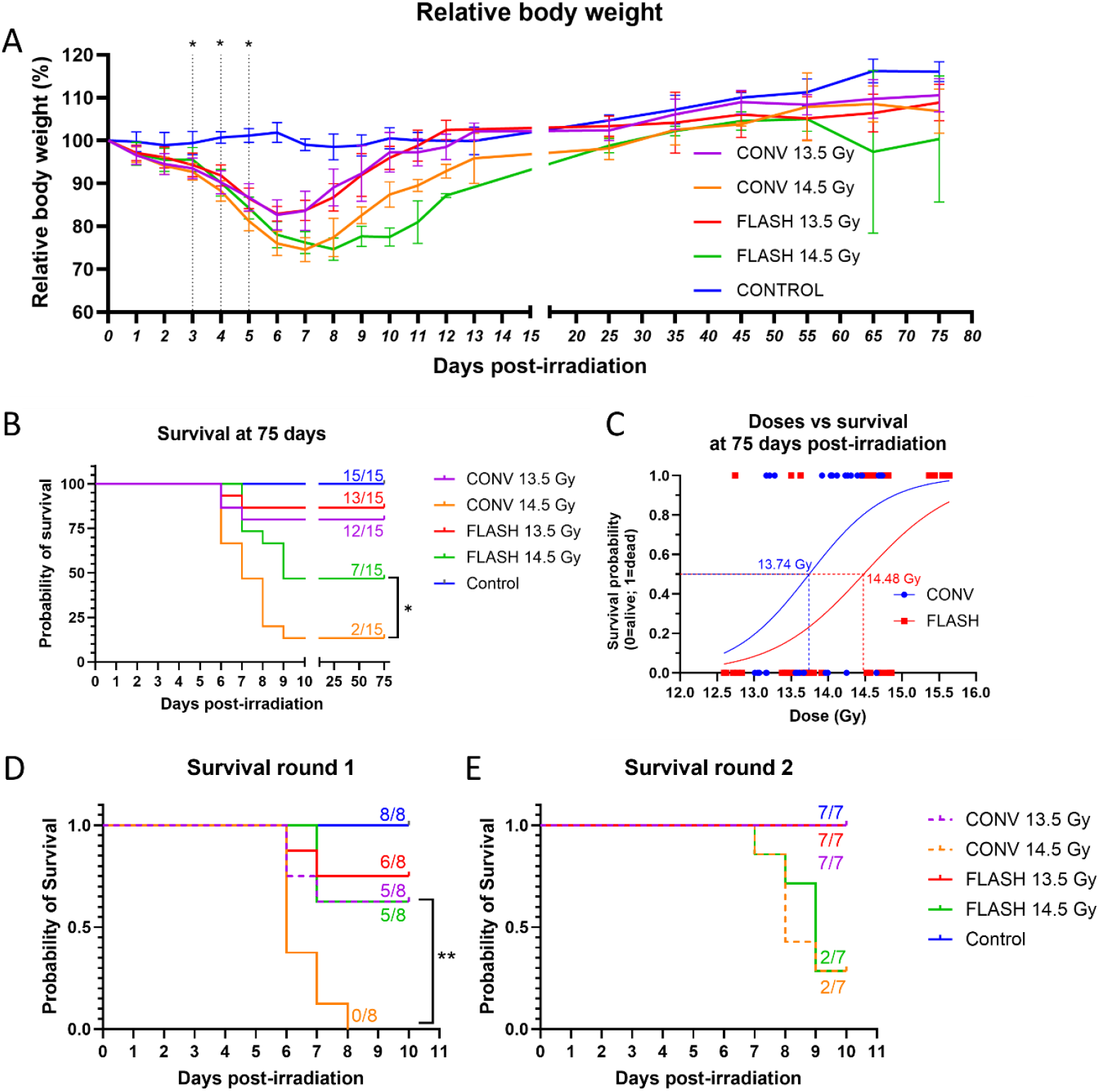
Weight loss and survival following FLASH and CONV proton irradiation. (A) Relative body weight (% of weight on the day of irradiation) of mice exposed to 13.5 Gy or 14.5 Gy of either FLASH (>60 Gy/s) or CONV (0.5 Gy/s) irradiation. A multiple t-test test was performed to test the statistical significance between the FLASH and CONV groups (*, p<0.05, FLASH 14.5 Gy vs. CONV 14.5 Gy); error bars indicate SD. (B) Kaplan-Meier survival plot of mice that received 13.5 or 14.5 Gy abdominal irradiation at FLASH or CONV dose rates. Events recorded mortality or euthanasia if mice demonstrated signs of severe morbidity, or if their weight decreased by 25% of the initial body weight. Survival curves were compared using the Log-rank (Mantel-Cox) test (*, p<0.05). (C) Simple logistic regression curve fit to the survivorship information against the alanine dose readings. Corresponding LD50 values were calculated for the FLASH and conventional cohorts. (D) Kaplan-Meier survival plot of mice irradiated during irradiation round 1. (E) Kaplan-Meier survival plot of mice irradiated during irradiation round 2.

**Figure 3.**
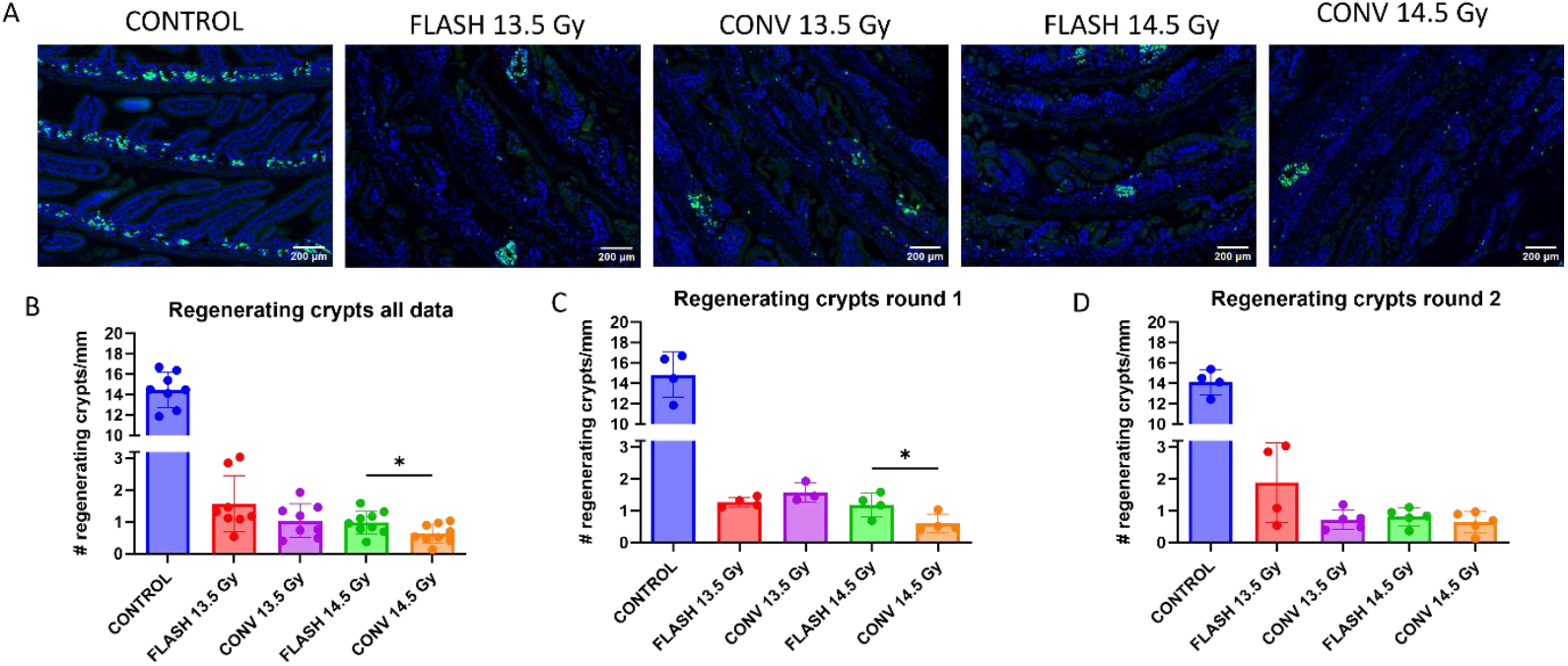
Regenerating intestinal crypts at four days following FLASH vs. CONV proton irradiation. (A) Representative images of EdU staining in intestinal sections at four days following 13.5 Gy and 14.5 Gy of abdominal irradiation. (B-D) Quantification of the average number of regenerating crypts per millimeter of intestine in mice (n=8-9 per condition) at four days following 13.5 Gy and 14.5 Gy abdominal proton irradiation at either FLASH or CONV dose rates: (B) pooled data, (C) data of irradiation round 1, (D) data of irradiation round 2. Regenerating crypts (≥5 EdU^+^ cells/crypt) were counted in two sections per mouse. A t-test was performed (*, p<0.05) to compare the effect of the same dose of CONV and FLASH treatment; error bars indicate SD.

### Increased regenerating intestinal crypt numbers following FLASH vs. CONV proton irradiation

To assess acute intestinal toxicity, crypt cell proliferation was measured by EdU labeling on day 4 post-irradiation (Figure 3). At 13.5 Gy, regenerating crypt numbers did not differ significantly between FLASH and CONV groups (1.57 vs. 1.04 crypts/mm). At 14.5 Gy, FLASH mice showed significantly more regenerating crypts than CONV (0.98 vs. 0.63 crypts/mm; p=0.04), consistent with survival outcomes (Figure 3B). Given the conflicting survival outcomes observed in two irradiation rounds, regenerating crypt numbers were also analyzed in individual batches. When analyzed per irradiation round, the sparing effect appeared at 14.5 Gy in round 1 (Figure 3C) (1.19 vs. 0.61 crypts/mm; p=0.05) but at 13.5 Gy in round 2 (1.88 vs. 0.72 crypts/mm; p=0.08). Overall, FLASH irradiation spared intestinal crypts in both experiments, though the effective dose shifted between rounds.

### Similar blood cell counts at four days following FLASH vs. CONV proton irradiation

To assess whether FLASH irradiation spares circulating immune cells, whole blood counts were performed on day 4 post-irradiation (Figure 4). Total white blood cells, lymphocytes, monocytes, and granulocytes were comparable between FLASH and CONV groups, with no significant differences in hematologic toxicity observed in pooled (Figure 4H) or round-specific analyses (Supplementary Figure 5).

**Figure 4.**
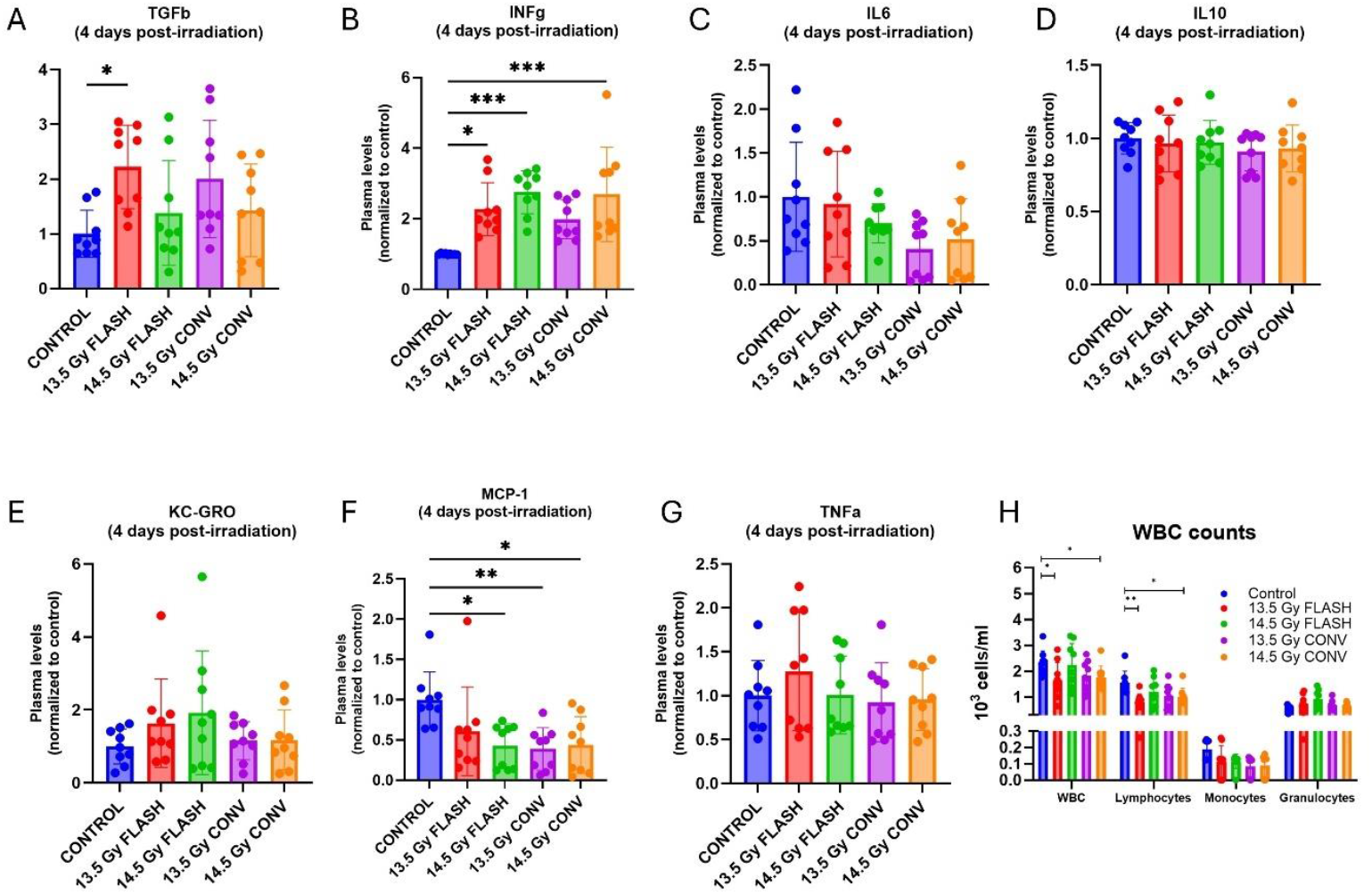
White blood cell counts and cytokine and chemokine expression in blood plasma 4 days following 13.5 Gy or 14.5 Gy of abdominal irradiation at FLASH or CONV dose rates. **(A-G)** The individual expression levels of TGFβ1, INFγ, IL6, IL10, KC-GRO/CXCL-1, MCP-1, and TNFα were normalized to the average of the control group levels of the same batch of animals (n=9 per condition). Differences between conditions were determined by a one-way ANOVA test with Tukey’s post hoc test (*, p<0.05; **, p<0.01, ***, p<0.001); error bars indicate SD. **(H)** Blood counts at four days following FLASH vs. CONV proton irradiation. Quantification of total white blood cell (WBC), lymphocytes, monocytes, and granulocytes in blood of mice (n=9 per condition) at four days following 13.5 Gy or 14.5 Gy of abdominal irradiation at FLASH or CONV dose rates. Differences between conditions were determined by a Kruskal-Wallis test with Dunn’s post hoc test (*, p<0.05; **, p<0.01); error bars indicate SD.

### Similar cytokine and chemokine levels at 4 and 75 days following FLASH vs. CONV proton irradiation

To assess potential differences in inflammatory responses after FLASH vs. CONV abdominal irradiation, five cytokines (TGFβ1, INFγ, IL6, IL10, TNFα) and two chemokines (KC-GRO/CXCL-1, MCP-1) were measured in mouse plasma at 4 and 75 days post-irradiation (Figures 4 and 5). At day 4, INFγ levels were consistently increased and MCP-1 decreased across all irradiated groups compared to controls, without significant differences between dose rates. At day 75, TGFβ1 tended to increase in all irradiated groups, reaching significance in 13.5 Gy FLASH, while TNFα was elevated in 13.5 Gy and 14.5 Gy FLASH and in 13.5 Gy CONV. Overall, cytokine and chemokine expression patterns were similar across doses and dose rates. The individual irradiation round results can be found in Supplementary Figures 7-10.

**Figure 5.**
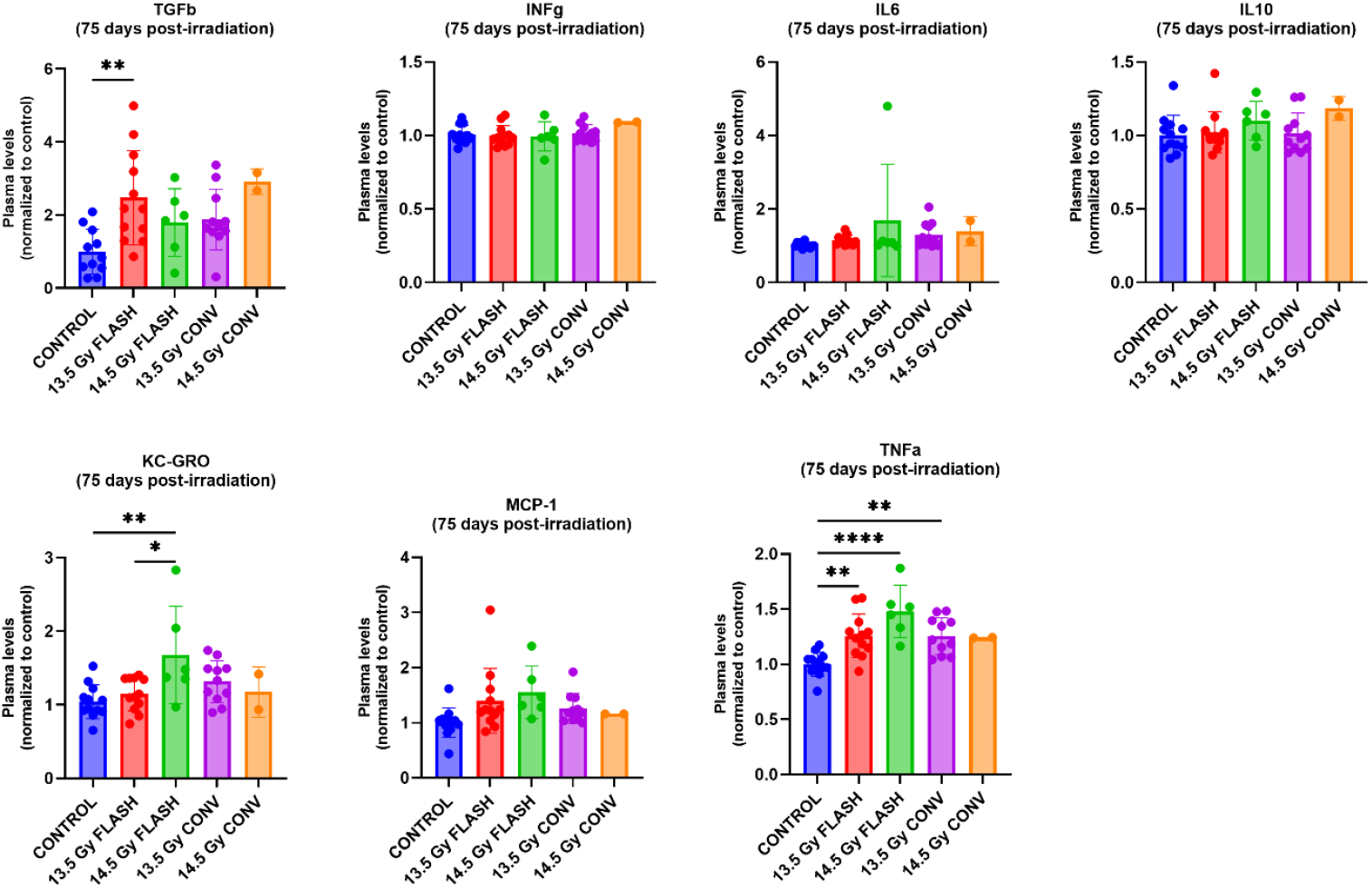
Cytokine and chemokine expression in blood plasma 75 days following 13.5 Gy or 14.5 Gy of abdominal irradiation at FLASH or CONV dose rates. The individual expression levels of TGFβ1, INFγ, IL6, IL10, KC-GRO/CXCL-1, MCP-1, and TNFα were normalized to the average of the control group levels of the same batch of animals (n=2-12 per condition). Differences between conditions were determined by a one-way ANOVA test with Tukey’s post hoc test (*, p<0.05; **, p<0.01, ***, p<0.001); error bars indicate SD.

### CONV and FLASH proton irradiation result in comparable structural intestinal changes at 75 days post-irradiation

To assess long-term effects, intestinal wall thickness was measured at 75 days post-irradiation as a marker of fibrosis (Figure 6). All treatments except 14.5 Gy FLASH caused significant thickening, with no overall differences between irradiated groups. Interestingly, in irradiation round 1, FLASH irradiation at 13.5 Gy caused a non-significant thickening of the muscle layer (1.06-fold vs. control), while the same dose of CONV irradiation led to a pronounced and statistically significant increase in intestinal wall thickness (1.22-fold vs. control), comparable to the effect of 14.5 Gy FLASH (1.17-fold vs. control) (Figure 5). Masson’s trichrome staining confirmed greater collagen deposition in CONV 13.5 Gy and FLASH 14.5 Gy, whereas FLASH 13.5 Gy samples were closer to non-irradiated tissue. No such differences were seen in round 2, consistent with early toxicity outcomes. In addition, cyst-like structures resembling early micro-adenomas [39] were observed in all irradiated groups, with lower incidence in 13.5 Gy FLASH, though not significantly different from other treatments (Figure 6C and Supplementary Figure 6 for individual irradiation rounds results).

**Figure 6.**
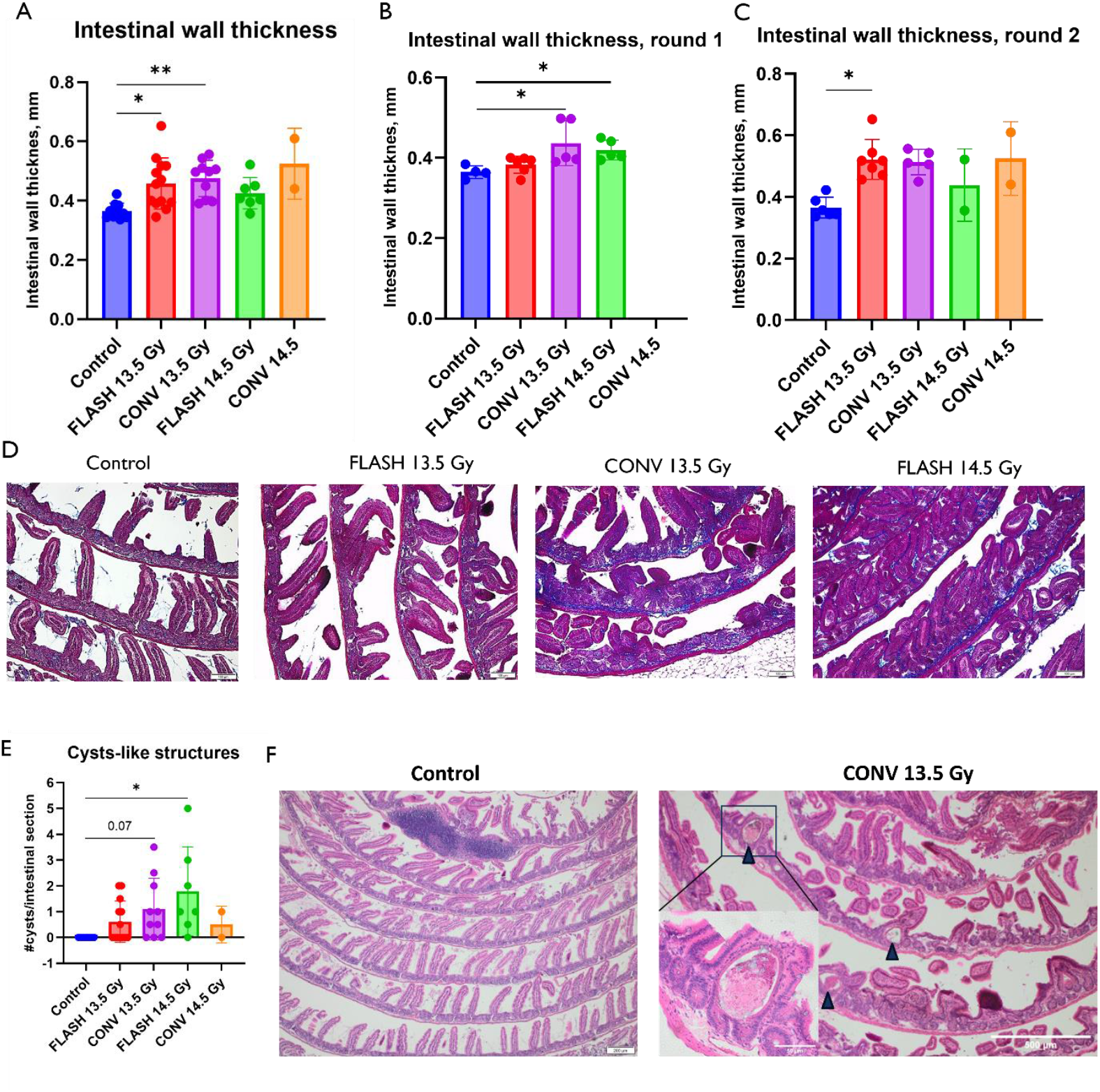
Structural changes in the small intestine at 75 days following FLASH vs. CONV irradiation. (A) Quantification of the mean thickness of the intestinal wall of mice (n=2-13 per condition) at 75 days following FLASH and CONV irradiation. The mean thickness was quantified by measuring the total intestinal wall thickness at 40-80 different locations within two non-adjacent intestinal sections per sample. (B) Mean thickness of the intestinal wall of mice irradiated during irradiation round 1. (C) Mean thickness of the intestinal wall of mice irradiated during irradiation round 2. (D) Masson’s Trichrome staining images of the murine intestine at 75 days following FLASH and CONV irradiation (from round 1) to visualize collagen deposition (blue staining). (E) Quantification of the number of cyst-like structures in the intestine of mice (n=2-13 per condition) at 75 days following FLASH and CONV irradiation. (F) H&E images of intestinal samples of non-irradiated controls and CONV-irradiated mice at 75 days post-irradiation showing multiple cyst-like structures (arrowheads). A Kruskal-Wallis test with Dunn’s post hoc test was performed (*, p<0.05, **, p<0.01); error bars indicate SD. De

## Discussion

In this proof-of-concept study, our goal was to demonstrate the feasibility and the intestinal tissue sparing effect of proton FLASH irradiation using a pulsed pencil beam generated by a superconducting synchrocyclotron.

We first assessed the feasibility of synchrocyclotron-based proton FLASH using PBS, the current clinical standard for its superior normal tissue sparing [26] and the most attractive system for clinical FLASH implementation. With this irradiation approach, we achieved stable and reproducible mean dose rates exceeding 60 Gy/s, a level considered sufficient to elicit the FLASH effect.

Second, the potential benefits of proton FLASH irradiation were evaluated in a mouse model of intestinal toxicity. At 14.5 Gy, FLASH significantly improved survival compared to CONV irradiation, with 87% and 53% mortality over the two irradiation rounds in the CONV and FLASH group, respectively. A FLASH modifying factor of 0.95 was obtained. Even though these overall results are promising and confirm that FLASH irradiation on a clinical synchrocyclotron is able to reduce intestinal toxicity in a mouse model, the obvious discrepancies between the two irradiation rounds cannot be ignored. Previously published *in vivo* data demonstrated improved survival after FLASH abdominal irradiation with electrons [27, 28], photons [29, 30] and scattered protons [31, 32]. However, three studies [33-35] showed either no benefit or worse outcomes, despite comparable or higher dose rates. Our data show that conflicting results in this particular model are observed not only by different research groups using different irradiation machines with different dose rates, doses, beam structure, mouse strain, anesthesia, irradiation field size, experimental endpoints and animal follow-up approaches to name a few. In our study, all experimental procedures were standardized and performed by the same investigators, with no variation in physical beam parameters or any other controllable conditions. Nevertheless, the improved survival observed after FLASH irradiation in our initial experiments could not be reproduced in the second round of irradiations.

To determine the acute effects of proton FLASH irradiation on the intestinal tissue, regeneration of intestinal crypts was evaluated at four days post-irradiation. Several electron FLASH studies demonstrated a 2-3-fold increase in regenerating crypts at four days following 11.2-12.5 Gy [36], 14 Gy [28], or 16 Gy [37] of FLASH vs. CONV irradiation. In addition, proton FLASH irradiation has shown its capacity in increasing the number of proliferating cells per crypt and the percentage of regenerating crypts in comparison to CONV irradiation, as measured by EdU staining [14]. Conversely, Bell and co-authors [33] and Zhang and co-authors [34] observed no differences in intestinal crypt cell proliferation and survival following FLASH and conventional dose rate proton irradiation. Finally, a recent study reported increased manifestation of early RT-induced GI toxicity (survival and crypt regeneration) after synchrotron-based proton FLASH versus CONV, but not after linac-based electron FLASH [35]. In our case, the intestinal crypt FLASH sparing effect shifted from 14.5 Gy in first irradiation round to 13.5 Gy in second irradiation round, an observation clearly requiring further investigation. On average, a significant increase in the number of regenerating crypts was induced by 14.5 Gy FLASH compared to 14.5 Gy CONV treatment, while no differences were observed after 13.5 Gy irradiation, corroborating our survival data.

To explore immune contributions, blood counts at day 4 showed no difference between FLASH and CONV, aligning with previous studies [12], [30], [33]. No *in vivo* study to date has demonstrated convincing lymphocyte sparing, and computational models suggest such effects are only expected at higher doses (≥20 Gy) and smaller fields [38, 39].

At day 75, intestinal wall thickening indicated fibrosis in all irradiated mice, with a clear trend toward reduced fibrosis in the FLASH 14.5 Gy group. The lack of significant long-term FLASH tissue sparing might possibly be explained by the relatively early tissue collection timepoint used in the current study, however, similar results were reported by Bell et al. who examined the expansion of lamina propria and submucosa layer at 8 months following 13 Gy FLASH and CONV irradiation [33], and Zhang et al. who examined thickening of muscular externa 280 days post-CONV vs. FLASH treatment with 16.2 Gy [34]. Diffenderfer et al., in contrast, showed that FLASH proton irradiation elicited significantly reduced intestinal fibrosis (thinner intestinal muscle layer) compared with CONV treatment at eight weeks following 18 Gy of local intestinal irradiation [14]. Cytokine analysis at day 75 showed no FLASH-specific reduction in IL6 or TGFβ1 levels, despite previous reports of modulation following abdominal [30] or lung and skin irradiation [20, 40, 41]. In this study, a trend of TGFβ1 increase was observed in all irradiated groups at both early and late timepoint, but significant differences between FLASH -and CONV-irradiated animals could not be found.

Taken together, our study confirms that FLASH proton irradiation delivered with a clinical synchrocyclotron PBS system can reduce acute intestinal toxicity in mice. However, the benefit is small, context-dependent, and not consistently reproducible. The extreme steepness of the abdominal irradiation dose–response curve in mice and the modest magnitude of the FLASH effect likely contribute to this variability. In fact, discrepancies in observed results in this model exist also beyond the FLASH field. For example, Brodin et al. report no deaths at doses below 16 Gy and 100% mortality at above 17 Gy following whole-abdominal X-ray irradiation [42], while Booth et al. showed that mice started dying at doses as low as 12 Gy and that 16 Gy resulted in 100% mortality [43]. Moreover, seemingly subtle differences in other factors might potentially have a significant impact on survival [44].

In conclusion, while our results support the technical feasibility of synchrocyclotron-based FLASH, they also highlight the limitations of murine GI toxicity as a preclinical model and emphasize the need for standardized protocols and mechanistic studies to resolve ongoing controversies.

## Declaration of author contributions

**Eva Bogaerts:** Conceptualization, methodology, formal analysis, investigation, visualization, original draft writing, review and editing. **Ellina Macaeva**: Conceptualization, methodology, project administration, supervision, formal analysis, investigation, visualization, original draft writing, review and editing. **Sima Qamhiyeh**: Methodology, investigation, review and editing. **Laurence Delombaerde**: Methodology, investigation, formal analysis, review and editing. **Brigitte Reniers**: Methodology, investigation, review and editing. **Marco Caprioli**: Methodology, investigation, review and editing. **Nicolas Gerard**: Methodology, investigation, formal analysis, review and editing. **Jarrick Nys**: Methodology, investigation, formal analysis, original draft writing, review and editing. **Valentin Hamaide**: Methodology, investigation, formal analysis, original draft writing, review and editing. **Alexis Warnier**: Methodology, investigation, review and editing. **Rudi Labarbe**: Conceptualization, methodology, review and editing. **Swati Girdhani**: conceptualization, methodology, review and editing. **Richard Coos**: methodology, review and editing. **Stéphane Lucas**: methodology, funding acquisition, review and editing. **Sofie Isebaert**: conceptualization, project administration, review and editing. **Rüveyda Dok**: methodology, supervision, review and editing. **Evelien Dierick**: project administration, review and editing. **Karin Haustermans**: conceptualization, supervision, review and editing. **Edmond Sterpin**: conceptualization, funding acquisition, supervision, review and editing

## Declaration of generative-AI and AI-assisted technologies in the writing process

During the preparation of this work the authors used ChatGPT to improve readability and language. After using this tool, the authors reviewed and edited the content as needed and take full responsibility for the content of the publication.

## References

1. Favaudon, V., et al., Ultrahigh dose-rate FLASH irradiation increases the differential response between normal and tumor tissue in mice. Sci Transl Med, 2014. 6(245): p. 245ra93.

2. Montay-Gruel, P., et al., Irradiation in a flash: Unique sparing of memory in mice after whole brain irradiation with dose rates above 100Gy/s. Radiother Oncol, 2017. 124(3): p. 365–369.

3. Tinganelli, W., et al., FLASH with carbon ions: Tumor control, normal tissue sparing, and distal metastasis in a mouse osteosarcoma model. Radiotherapy and Oncology, 2022.

4. Kim, M.M., et al., Comparison of FLASH Proton Entrance and the Spread-Out Bragg Peak Dose Regions in the Sparing of Mouse Intestinal Crypts and in a Pancreatic Tumor Model. Cancers (Basel), 2021. 13(16).

5. Gao, F., et al., First demonstration of the FLASH effect with ultrahigh dose-rate high energy X-rays. bioRxiv, 2020: p. 2020.11.27.401869.

6. Vozenin, M.C., et al., The Advantage of FLASH Radiotherapy Confirmed in Mini-pig and Cat-cancer Patients. Clin Cancer Res, 2019. 25(1): p. 35–42.

7. Iturri, L., et al., Proton FLASH Radiation Therapy and Immune Infiltration: Evaluation in an Orthotopic Glioma Rat Model. Int J Radiat Oncol Biol Phys, 2023. 116(3): p. 655–665.

8. Ghannam, Y., et al., First evidence of in vivo effect of FLASH radiotherapy with helium ions in zebrafish embryos. Radiother Oncol, 2023. 187: p. 109820.

9. Mascia, A.E., et al., Proton FLASH Radiotherapy for the Treatment of Symptomatic Bone Metastases: The FAST-01 Nonrandomized Trial. JAMA Oncol, 2023. 9(1): p. 62–69.

10. Daugherty, E.C., et al., FLASH radiotherapy for the treatment of symptomatic bone metastases in the thorax (FAST-02): protocol for a prospective study of a novel radiotherapy approach. Radiat Oncol, 2024. 19(1): p. 34.

11. Bell, B.I., et al., Whole Abdominal Pencil Beam Scanned Proton FLASH Increases Acute Lethality. International Journal of Radiation Oncology, Biology, Physics.

12. Zhang, Q., et al., Absence of Tissue-Sparing Effects in Partial Proton FLASH Irradiation in Murine Intestine. Cancers, 2023. 15(8): p. 2269.

13. Ribeiro, C.O., et al., Evaluation of continuous beam rescanning versus pulsed beam in pencil beam scanned proton therapy for lung tumours. Physics in Medicine & Biology, 2020. 65(23): p. 23NT01.

14. Diffenderfer, E.S., et al., Design, Implementation, and in Vivo Validation of a Novel Proton FLASH Radiation Therapy System. Int J Radiat Oncol Biol Phys, 2020. 106(2): p. 440–448.

15. Zhang, Q., et al., FLASH Investigations Using Protons: Design of Delivery System, Preclinical Setup and Confirmation of FLASH Effect with Protons in Animal Systems. Radiat Res, 2020. 194(6): p. 656–664.

16. Girdhani, S., et al., Abstract LB-280: FLASH: A novel paradigm changing tumor irradiation platform that enhances therapeutic ratio by reducing normal tissue toxicity and activating immune pathways. Cancer Research, 2019. 79(13_Supplement): p. LB-280-LB-280.

17. Beyreuther, E., et al., Feasibility of proton FLASH effect tested by zebrafish embryo irradiation. Radiother Oncol, 2019. 139: p. 46–50.

18. Sørensen, B.S., et al., Pencil beam scanning proton FLASH maintains tumor control while normal tissue damage is reduced in a mouse model. Radiotherapy and Oncology, 2022. 175: p. 178–184.

19. Leavitt, R.J., et al., Acute Hypoxia Does Not Alter Tumor Sensitivity to FLASH Radiation Therapy. Int J Radiat Oncol Biol Phys, 2024. 119(5): p. 1493–1505.

20. Cunningham, S., et al., FLASH Proton Pencil Beam Scanning Irradiation Minimizes Radiation-Induced Leg Contracture and Skin Toxicity in Mice. Cancers, 2021. 13(5): p. 1012.

21. W. Kleeven, M.A.E. Forton, S. Henrotin, Y. Jongen, V. Nuttens, Y. Paradis, E. Pearson, S. Quets, J. Van de Walle, P. Verbruggen, S. Zaremba. THE IBA SUPERCONDUCTING SYNCHROCYCLOTRON PROJECT S2C2. in Proceedings of Cyclotrons 2013. 2013. Vancouver, BC, Canada.

22. Crijns, W., et al., Calibrating page sized Gafchromic EBT3 films. Med Phys, 2013. 40(1): p. 012102.

23. Silvestre Patallo, I.A.R; Coos, R; Hussein, M; Grimwood, A; Lucas, S; Hawkins, M; Baker, C; Palmans, H; Nisbet, A, Preliminary dosimetric validation of a small animal irradiation box for preclinical irradiation research in clinical proton beam centres, in 61st Annual Conference of the Particle Therapy Cooperative Group (PTCOG 61). 2023: Marid, Spain.

24. Bialkowska, A.B., et al., Improved Swiss-rolling Technique for Intestinal Tissue Preparation for Immunohistochemical and Immunofluorescent Analyses. J Vis Exp, 2016(113).

25. Deffet, S., V. Hamaide, and E. Sterpin, Definition of dose rate for FLASH pencil-beam scanning proton therapy: A comparative study. Med Phys, 2023. 50(9): p. 5784–5792.

26. Chuong, M., et al., Pencil beam scanning versus passively scattered proton therapy for unresectable pancreatic cancer. J Gastrointest Oncol, 2018. 9(4): p. 687–693.

27. Loo, B.W., et al., (P003) Delivery of Ultra-Rapid Flash Radiation Therapy and Demonstration of Normal Tissue Sparing After Abdominal Irradiation of Mice. International Journal of Radiation Oncology, Biology, Physics, 2017. 98(2): p. E16.

28. Levy, K., et al., Abdominal FLASH irradiation reduces radiation-induced gastrointestinal toxicity for the treatment of ovarian cancer in mice. Scientific Reports, 2020. 10(1): p. 21600.

29. Gao, F., et al., First demonstration of the FLASH effect with ultrahigh dose rate high-energy X-rays. Radiother Oncol, 2022. 166: p. 44–50.

30. Zhu, H., et al., Radioprotective effect of X-ray abdominal FLASH irradiation: Adaptation to oxidative damage and inflammatory response may be benefiting factors. Med Phys, 2022. 49(7): p. 4812–4822.

31. Evans, T., et al., Demonstration of the FLASH Effect Within the Spread-out Bragg Peak After Abdominal Irradiation of Mice. Int J Part Ther, 2022. 8(4): p. 68–75.

32. Liu, K., et al., Redefining FLASH Radiation Therapy: The Impact of Mean Dose Rate and Dose Per Pulse in the Gastrointestinal Tract. Int J Radiat Oncol Biol Phys, 2025. 121(4): p. 1063–1076.

33. Bell, B.I., et al., Whole Abdominal Pencil Beam Scanned Proton FLASH Increases Acute Lethality. Int J Radiat Oncol Biol Phys, 2025. 121(2): p. 493–505.

34. Zhang, Q., et al., Absence of Tissue-Sparing Effects in Partial Proton FLASH Irradiation in Murine Intestine. Cancers (Basel), 2023. 15(8).

35. Liu, K., et al., Discordance in Acute Gastrointestinal Toxicity between Synchrotron-Based Proton and Linac-based Electron Ultra-High Dose Rate Irradiation. Int J Radiat Oncol Biol Phys, 2025. 122(2): p. 491–501.

36. Ruan, J.L., et al., Irradiation at Ultra-High (FLASH) Dose Rates Reduces Acute Normal Tissue Toxicity in the Mouse Gastrointestinal System. Int J Radiat Oncol Biol Phys, 2021. 111(5): p. 1250–1261.

37. Eggold, J.T., et al., Abdominopelvic FLASH Irradiation Improves PD-1 Immune Checkpoint Inhibition in Preclinical Models of Ovarian Cancer. Mol Cancer Ther, 2022. 21(2): p. 371–381.

38. Galts, A. and A. Hammi, FLASH radiotherapy sparing effect on the circulating lymphocytes in pencil beam scanning proton therapy: impact of hypofractionation and dose rate. Physics in Medicine & Biology, 2024. 69(2): p. 025006.

39. Jin, J.Y., et al., Ultra-high dose rate effect on circulating immune cells: A potential mechanism for FLASH effect? Radiother Oncol, 2020. 149: p. 55–62.

40. Favaudon, V., et al., Ultrahigh dose-rate FLASH irradiation increases the differential response between normal and tumor tissue in mice. Science Translational Medicine, 2014. 6(245): p. 245ra93–245ra93.

41. Velalopoulou, A., et al., FLASH Proton Radiotherapy Spares Normal Epithelial and Mesenchymal Tissues While Preserving Sarcoma Response. Cancer Res, 2021. 81(18): p. 4808–4821.

42. Brodin, N.P., et al., A Model for Precise and Uniform Pelvic- and Limb-Sparing Abdominal Irradiation to Study the Radiation-Induced Gastrointestinal Syndrome in Mice Using Small Animal Irradiation Systems. Dose Response, 2017. 15(1): p. 1559325816685798.

43. Booth, C., et al., Evidence of delayed gastrointestinal syndrome in high-dose irradiated mice. Health Phys, 2012. 103(4): p. 400–10.

44. DiCarlo, A.L., et al., Study logistics that can impact medical countermeasure efficacy testing in mouse models of radiation injury. Int J Radiat Biol, 2021. 97(Sup1): p. S151–S167.

