## Supplementary material for "FLASH or flare: variable intestinal toxicity results in a mouse model following proton pencil beam scanning irradiation on a clinical superconducting synchrocyclotron"

### Supplementary Materials and Methods

#### *S2C2 beam time structure*

The IBA S2C2 synchrocyclotron accelerates one pulse of protons every millisecond (Pulse Repetition Frequency, *PRF* = 1 kHz) [1]. As the pulse lasts for 7 μs ($t_{pulse}$), there are 993 μs between subsequent pulses ($\Delta t_{pulse}$). The beam intensity is modulated only by means of charge per pulse ($Q_{pulse}$) modification, which in turn is modulated through the average radiofrequency system amplitude during the whole acceleration cycle. Schematic representation of the S2C2 beam time structure is shown in Figure 1.

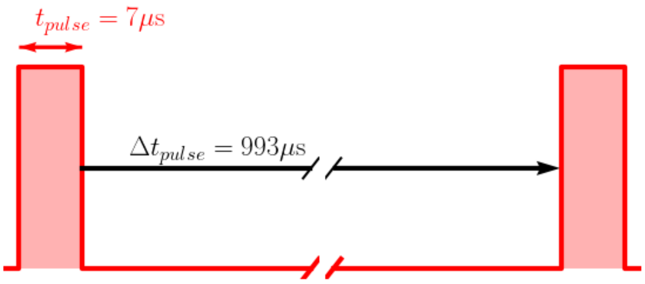


**Figure 1. Schematic representation of the S2C2 beam time structure.** The red line represents the instantaneous beam current extracted from the S2C2 ($Q_{pulse}$). The shaded red area is the pulse charge ($Q_{pulse}$).

#### *Treatment plan*

Pencil Beam Scanning technique was used to deliver the shoot through beam. The single-energy treatment plan was characterized by the mean beam energy (226.5 MeV for CONV and 229 MeV for UHDR), the total requested Monitor Units (MUs), the spot weights and lateral positions.

A custom-made brass aperture (20x20mm) was used to constrain the lateral profile. Both treatment plans thus had a larger field size than the aperture opening. Even though this slightly decreased the dose rate, it allowed to obtain a better dose homogeneity and robustness to possible aperture misalignment.

Two treatment plans were designed for CONV and UHDR irradiations. The CONV treatment plan consists of 49 spots of equal weights arranged in a square field of

27x27mm with a horizontal and vertical 4.5mm spot spacing. The UHDR treatment plan consisted of 25 spots arranged in a square field of 28x28mm with 7mm spot spacing, keeping the same ratio of spot spacing to spot size as the CONV case. In order to achieve ultra-high dose rates, the beam line optics were optimized to maximize the transmission from the S2C2 to isocenter, leading to an increased spot size at isocenter compared to the CONV treatment. The UHDR spot had a small tail towards the Y-direction (Figure 2). To achieve a uniform field, the spot weight was modulated in this axis.

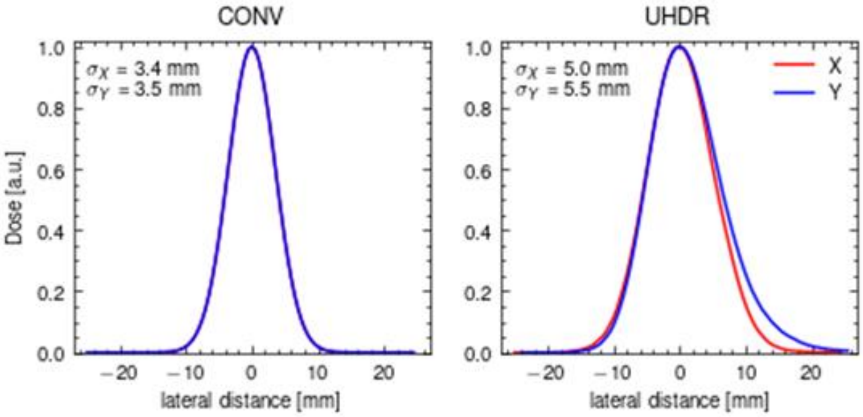


**Figure 2:** Spot shape for conventional PBS treatment (left) and ultra-high dose rate (right).

#### *Beam delivery and timings*

In the UHDR mode, each beam deflection was done in between pulses (<993μs). In between scannings, the beam was delivered in multiple pulses. The number of pulses per spot (see Table 1 for examples) depended on the requested MUs per spot ($MUPS_{REQ}$) for that spot and the maximum MUs per pulse available ($MUPP_{MAX}$):

$$\#_{\text{pulse}} = \left\lfloor \frac{\text{MUPS}_{\text{REQ}}}{\text{MUPP}_{\text{MAX}}} \right\rfloor \qquad (1)$$

Table 1.

| Target Dose [Gy] | Date (month-year) | $\#_{\text{pulse}}$ |
|---|---|---|
| **13.5** | 04-2024 | 210 |
| **14.5** | 04-2024 | 220 |
| **12.5** | 10-2024 | 205 |
| **13.5** | 10-2024 | 220 |
| **14.5** | 10-2024 | 235 |

| | | |
|---|---|---|
| **15.5** | 10-2024 | 245 |

***In silico***

An *in silico* model of the entire experimental setup (custom-made brass aperture, mice, animal positioning box) was developed to study dose profile across depth inside the mice via Monte Carlo (MC) simulations.

The *in silico* simulations were performed in the research treatment planning system, OpenTPS [2], which uses MCsquare [3] as dose engine for MC computations. The animal positioning box and the aperture were modeled as a voxelized geometry with known dimensions and material composition, and the mouse was modeled as pure water with a height of 1.5cm, representing the average height of the mice used for this experiment.

After computing the dose distribution across the mice and the full experimental setup, slices from the simulated 3D dose distribution at specific locations were compared to the Gafchromic film measurements and alanine pellet readings.

The *in silico* model was validated with the film (EBT-XD) and alanine pellet measurements. During the experiment on mice, the doses measured on the dosimetry devices below the mice were also compared to the ones from the simulation

***Animals***

Mice were housed in a conventional animal facility with 12 h dark/light cycles in individually ventilated cages (4-5 mice/cage) and were given free access to drinking water and food. Mice were given at least one week of acclimatization before the start of the irradiation. Following irradiation, mice were checked and weighed daily and ethically euthanized upon onset of severe morbidity including hunched posture, rough fur/lack of grooming, relative immobility, dehydration, or weight loss >25%.

***Animal positioning***

Reproducible positioning within each compartment was achieved by securing the mouse with two elastic bands at fixed locations, and taping hindfeet and tail. One day prior to the irradiation treatment, positioning reproducibility to within 1 mm was confirmed by micro-cone beam CT imaging of mice. The next day, mice were placed in the IRRAMICE box, which was loaded into a polymethyl methacrylate (PMMA) cradle registered to specified positions relative to the S2SC treatment head for FLASH and CONV irradiation.

***EdU staining***

Approximately 15-cm long segments of the intestine, beginning from the stomach and including the duodenum and the jejunum, were collected by the Swiss-roll technique. Collected intestinal segments were flushed with modified Bouin's fixative (50% ethanol/5% acetic acid in PBS). Using scissors, the intestinal lumen was cut open longitudinally along the mesenteric line and rinsed in PBS. Tweezers were used to roll the intestinal segment from the proximal end, with the luminal side up. The tissues were then fixed with 10% formalin overnight and stored in PBS for paraffin embedding.

Tissue sections were incubated with the Click-iT reaction cocktail for 30 min, followed by washing steps in 3% BSA and 1X PBS. To visualize nuclei, sections were incubated with Hoechst 33342 for 30 min. Sections were mounted and imaged at 10X magnification on a BX43F microscope (Olympus). From each sample, the most severely damaged area, as defined by >3 mm region with least number of crypts, was selected as the region of interest, and the number of regenerating crypts (≥5 EdU+ cells/crypt) was counted.

**Histology**

The mean thickness of the intestinal wall (including the submucosa and muscularis propria), as a marker of intestinal fibrosis, was quantified by measuring the muscle layer at 40-80 different locations within two non-adjacent intestinal sections per sample using ImageJ software. In addition, the average number of intra-mucosal cyst-like structures was quantified in two non-adjacent intestinal sections per sample. To qualitatively check for collagen deposition, sections were stained with Masson’s trichrome using a Masson Trichrome kit (Biognost; #MST-K-500) following the manufacturer’s protocol. The sections were mounted and imaged at 10X magnification on a BX43F microscope (Olympus).

1. W. Kleeven, M.A., E. Forton, S. Henrotin, Y. Jongen, V. Nuttens, Y. Paradis, E. Pearson, S. Quets, J. Van de Walle, P. Verbruggen, S. Zaremba. *THE IBA SUPERCONDUCTING SYNCHROCYCLOTRON PROJECT S2C2*. in *Proceedings of Cyclotrons 2013*. 2013. Vancouver, BC, Canada.
2. S. Wuyckens, D.D., G. Janssens, V. Hamaide, M. Huet, E. Loÿen, G. Rotsart de Hertaing, B. Macq, E. Sterpin, J. A. Lee, K. Souris, S. Deffet, *OpenTPS -- Open-source treatment planning system for research in proton therapy*. 2023.
3. Souris, K., J.A. Lee, and E. Sterpin, *Fast multipurpose Monte Carlo simulation for proton therapy using multi- and many-core CPU architectures.* Med Phys, 2016. **43**(4): p. 1700.
