## Supplementary figures and tables for "FLASH or flare: variable intestinal toxicity results in a mouse model following proton pencil beam scanning irradiation on a clinical superconducting synchrocyclotron"

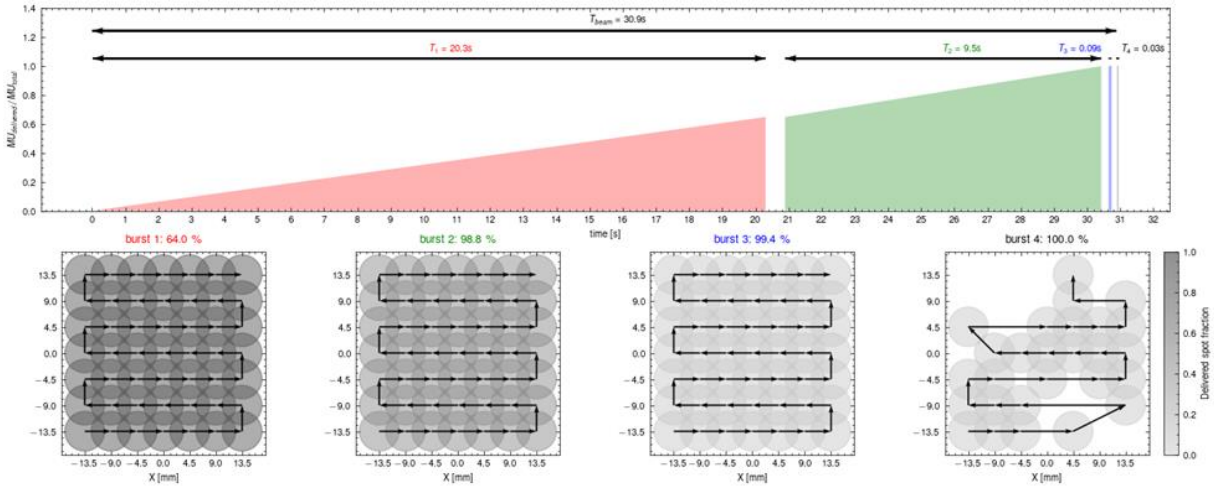


***Supplementary figure 1.*** *The CONV treatment plan is delivered in 4 bursts. Each sequential bursts delivers a smaller fraction of the dose at lower intensity that the burst before. (Top) The cumulative delivered monitor units in function of time for each of the bursts. The total treatment time (30.9s) is the sum of the delivery times of each of the bursts plus the time between bursts. (Bottom) The scanning pattern for each burst. In the final burst, some spots have already reached the requested monitor units, so they are not anymore part of the scanning pattern.*

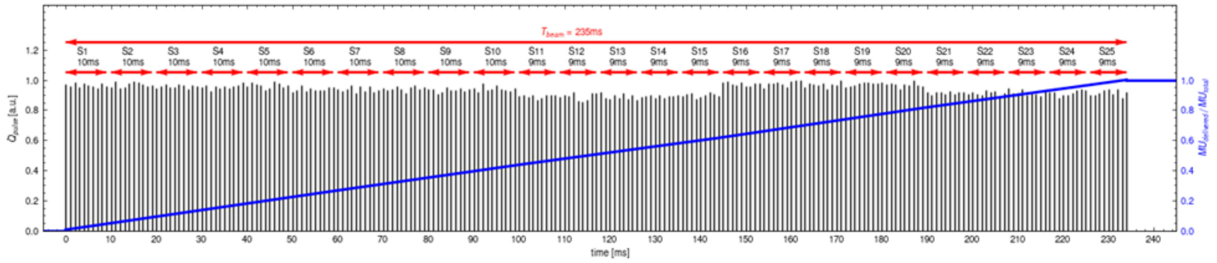


***Supplementary figure 2****1. Instantaneous current in function of time over the whole irradiation of a 14Gy UHDR irradiation. Each spike (black) represents a 7 µs pulse. Different spots are delivered with different number of pulses (See equation* ***Error! Reference source not found.****), depending on the requested monitor units. The total treatment time is 235ms. The delivered monitor units are nearly linear as a function of time.*

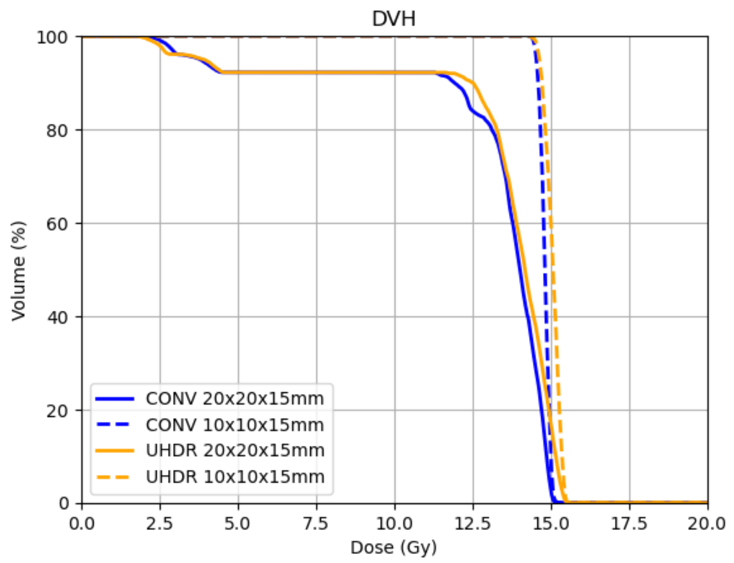


***Supplementary figure 3***. *Dose-volume histogram for the CONV and UHDR experiments at 14.5Gy on the whole target volume of size 20x20x15mm and the central subvolume of size 10x10x15mm.*

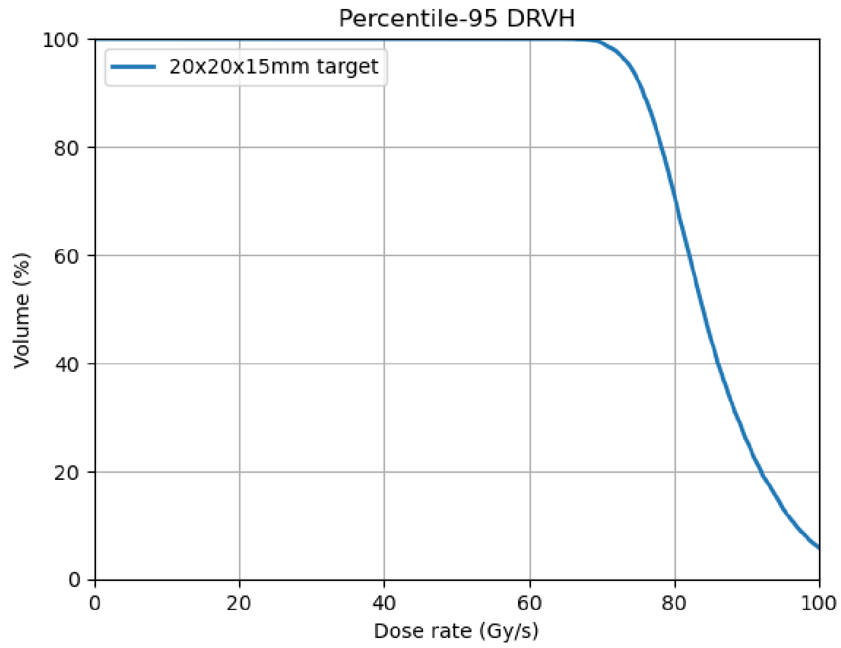


***Supplementary figure 4.*** *2 Dose rate volume histogram for the UHDR experiment at 14.5Gy on the 20x20x15mm target. The dose rate definition used in the max-percentile dose rate with a percentile = 95% (i.e. the local dose rate at the voxel level where the time is computed as the elapsed time to irradiate 95% of the received dose at that voxel). DR95=74Gy/s.*

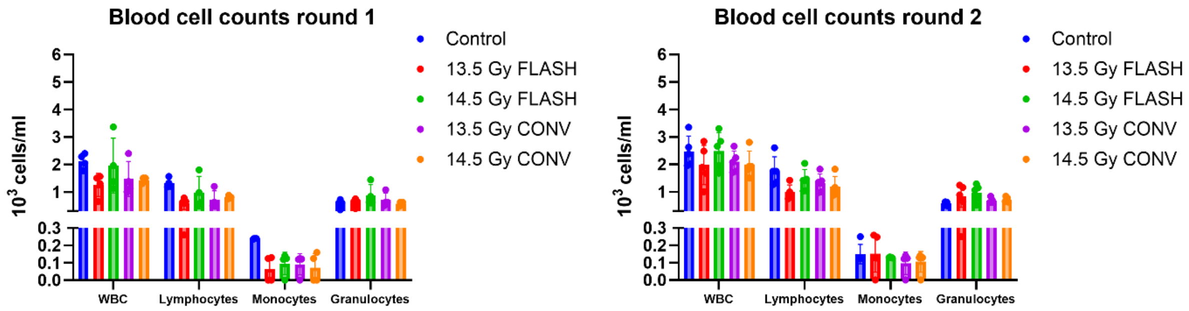


**Supplementary figure 5.** Blood counts at four days following FLASH vs. CONV proton irradiation in round 1 and 2.

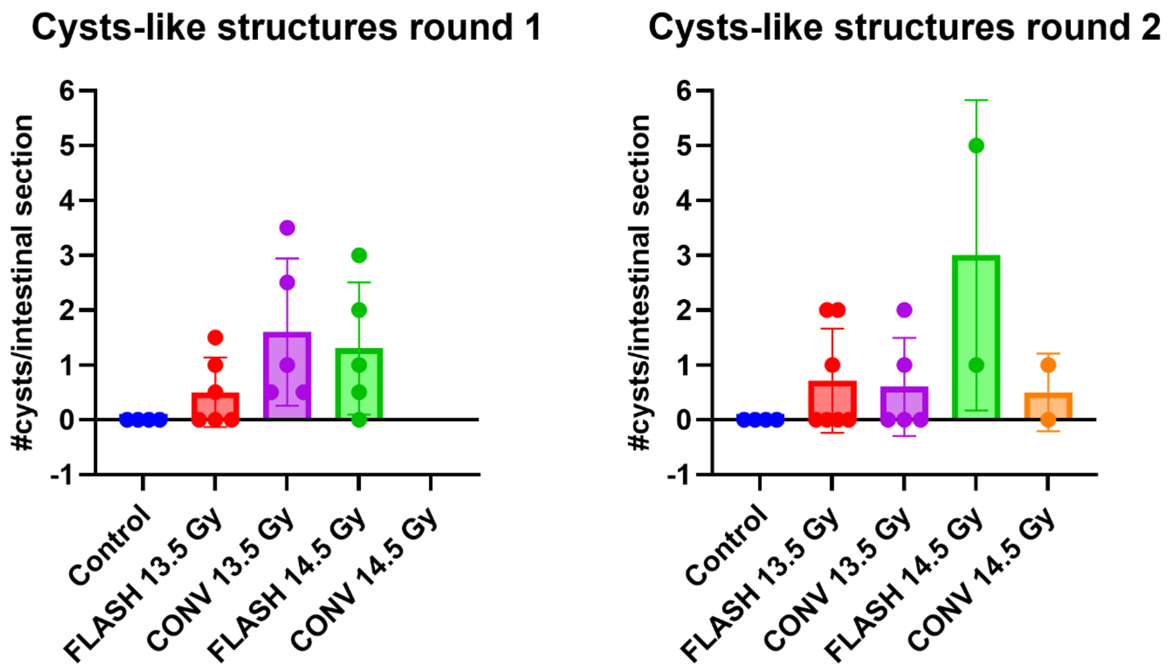


**Supplementary figure 6.** Cyst-like structure counts in the intestine of mice at 75 days following FLASH vs. CONV irradiation in round 1 and 2.

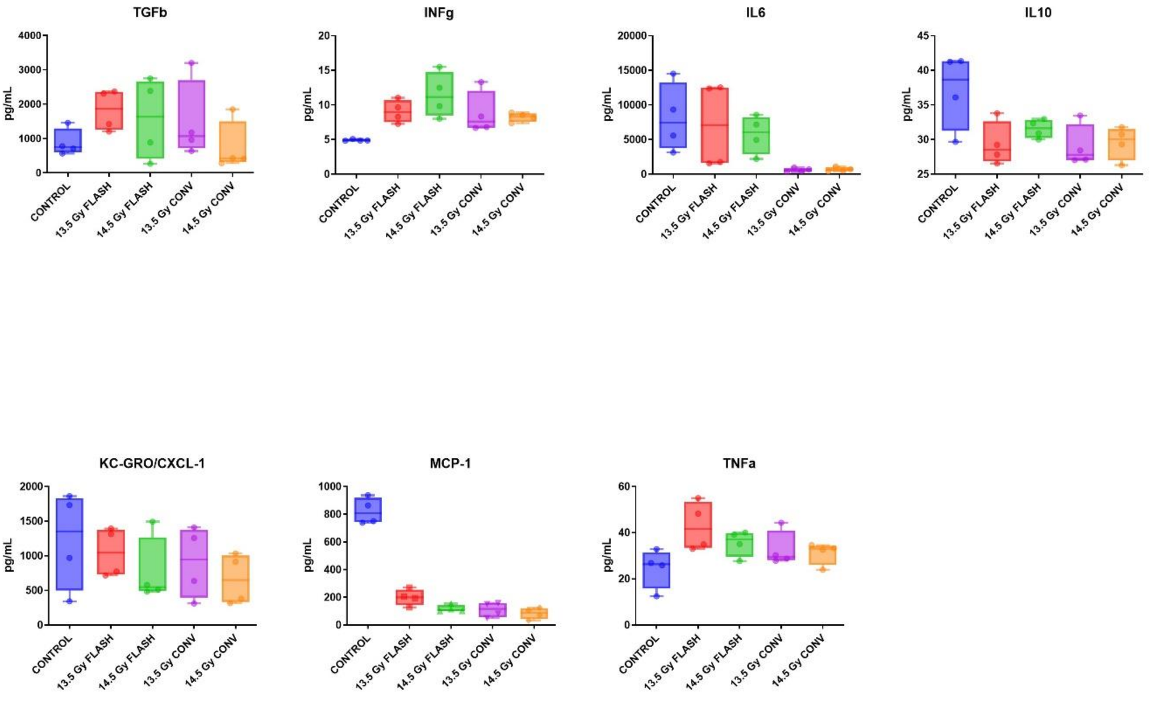


**Supplementary figure 7. Cytokine and chemokine expression in blood plasma 4 days following 13.5 Gy or 14.5 Gy of abdominal irradiation at FLASH or CONV dose rates in irradiation round 1.**

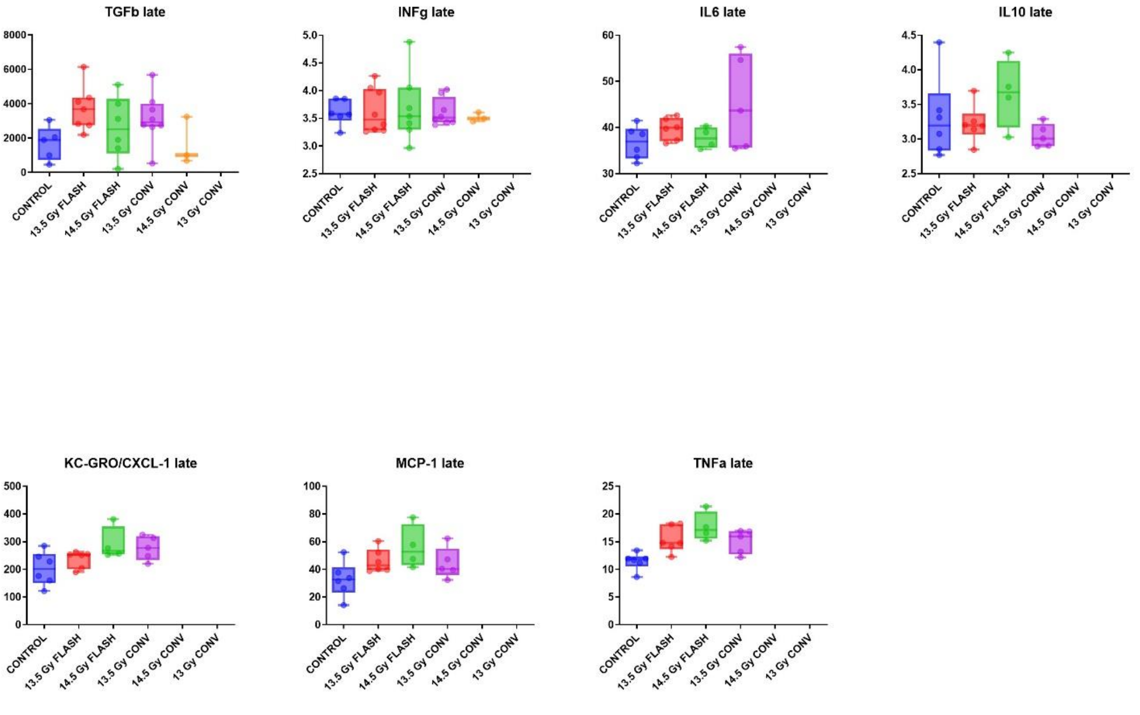


**Supplementary figure 8. Cytokine and chemokine expression in blood plasma 75 days following 13.5 Gy or 14.5 Gy of abdominal irradiation at FLASH or CONV dose rates in irradiation round 1.**

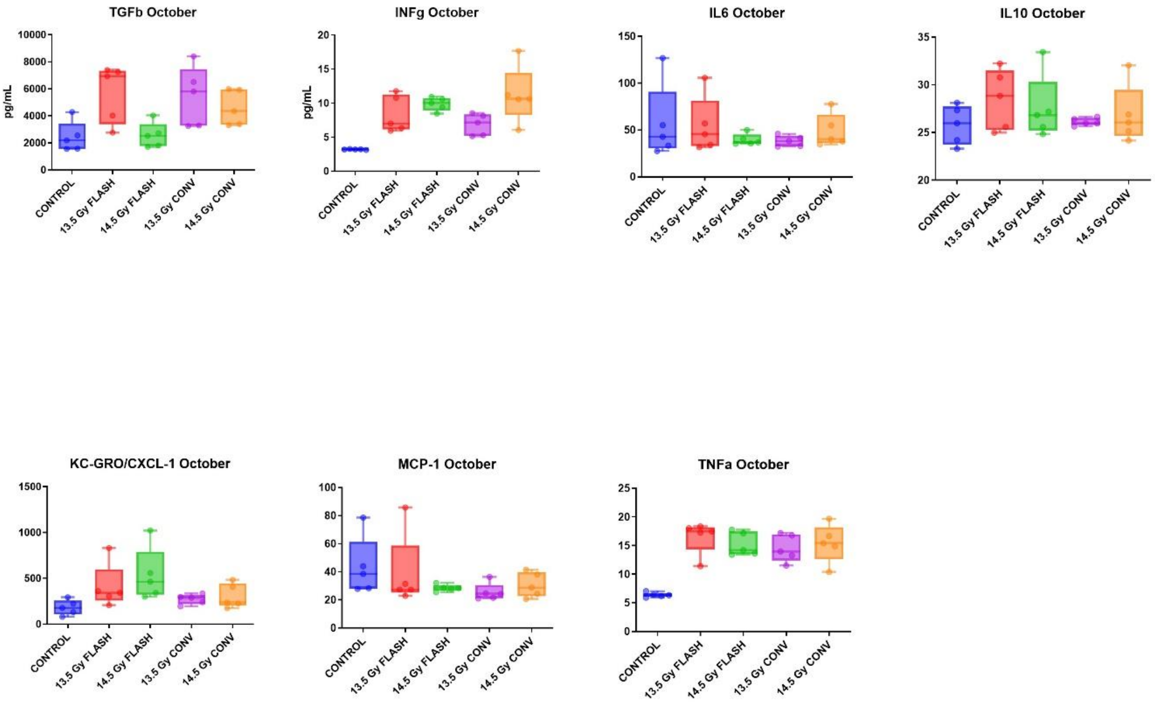


**Supplementary figure 9. Cytokine and chemokine expression in blood plasma 4 days following 13.5 Gy or 14.5 Gy of abdominal irradiation at FLASH or CONV dose rates in irradiation round 2.**

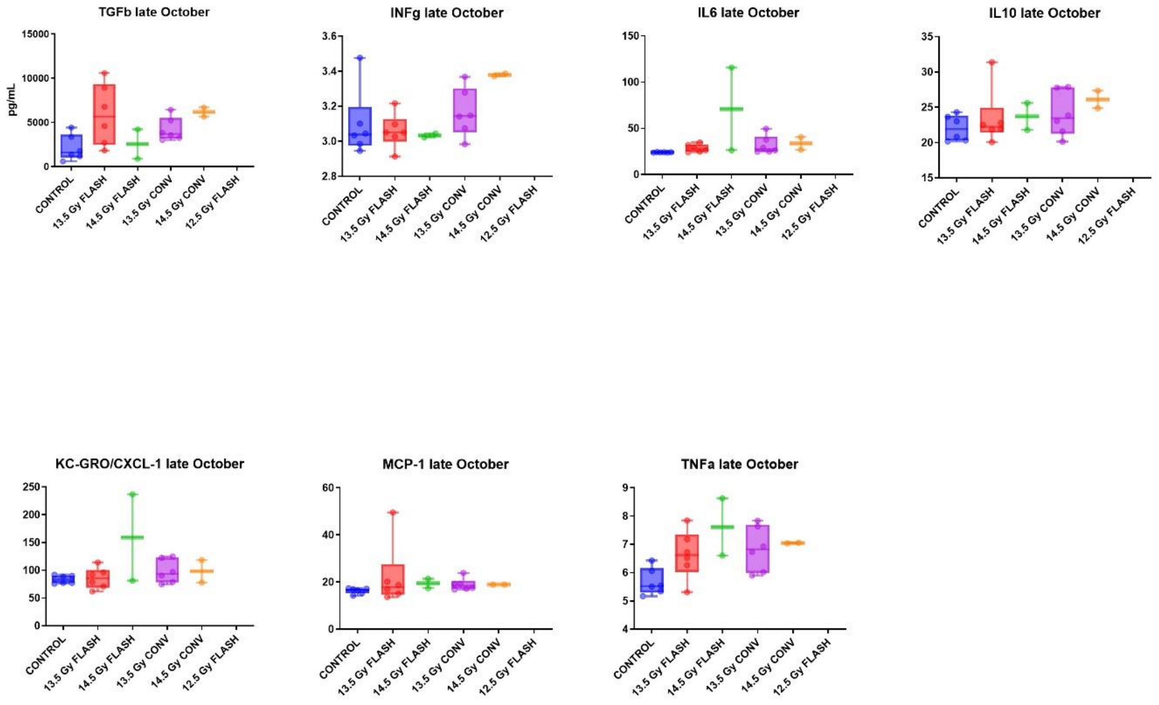


**Supplementary figure 10. Cytokine and chemokine expression in blood plasma 75 days following 13.5 Gy or 14.5 Gy of abdominal irradiation at FLASH or CONV dose rates in irradiation round 2.**

Supplementary tables

**Supplementary table 1.** Dosimetry results of film measurements on a specific batch of mice irradiated at 14.5Gy at conventional dose rate. The comparison is done for different region sizes.

| | Median dose on 20x20mm field at location of film | Median dose on 10x10mm field at location of film | Median dose on 5x5mm field at location of film | Alanine measure |
|---|---|---|---|---|
| Mouse 1 | 12.77 | 13.74 | 13.92 | 14.25 |
| Mouse 2 | 13.06 | 13.95 | 14.15 | 14.26 |
| Mouse 3 | 12.67 | 13.55 | 13.79 | 14.31 |
| Mouse 4 | 12.76 | 13.69 | 13.78 | 14.04 |
| Mouse 5 | 13.13 | 13.98 | 14.12 | 14.07 |

| | | | | |
|---|---|---|---|---|
| Mouse 6 | 12.86 | 13.79 | 13.97 | 14.15 |
| **Average** | **12.88** | **13.78** | **13.96** | **14.18** |
| **MC simu** | **13.69** | **14.24** | **14.37** | **14.25** |
| **Variation [%]** | **6.13%** | **3.26%** | **2.93%** | **0.49%** |

**Supplementary table 2.** Dosimetry results of film measurements on a specific batch of mice irradiated at 14.5Gy at UHDR dose rate. The comparison is done for different region sizes.

| | Median dose on 20x20mm field at location of film | Median dose on 10x10mm field at location of film | Median dose on 5x5mm field at location of film | Alanine measure |
|---|---|---|---|---|
| Mouse 1 | 12.84 | 13.81 | 14.03 | 14.42 |
| Mouse 2 | 12.98 | 13.82 | 14.00 | 14.41 |
| Mouse 3 | 13.01 | 13.99 | 14.30 | 14.31 |
| Mouse 4 | 12.83 | 13.83 | 14.14 | 14.26 |
| Mouse 5 | 13.24 | 14.12 | 14.30 | 14.28 |
| Mouse 6 | 12.96 | 13.94 | 14.15 | 14.10 |
| **Average** | **12.98** | **13.91** | **14.15** | **14.30** |
| **MC simu** | **13.76** | **14.29** | **14.38** | **14.22** |
| **Variation [%]** | **5.86%** | **2.63%** | **1.58%** | **0.54%** |
